# TMEM106B is increased in Multiple Sclerosis plaques, and deletion causes accumulation of lipid after demyelination

**DOI:** 10.1101/2022.05.05.490697

**Authors:** Bridget Shafit-Zagardo, Simone Sidoli, James E. Goldman, Juwen C. DuBois, John R. Corboy, Stephen M. Strittmatter, Hillary Guzik, Sarah Graff, Rashed M. Nagra

**Affiliations:** Department of Pathology, Albert Einstein College of Medicine, 1300 Morris Park Avenue Bronx, New York 10461, USA.; (senior author); Department of Biochemistry, Albert Einstein College of Medicine, 1300 Morris Park Avenue Bronx, New York 10461, USA.; Department of Pathology and Cell Biology, Columbia University College of Physicians and Surgeons. New York, NY; Department of Pathology, 1300 Morris Park Avenue Bronx, New York 10461, USA. 718430-2189; Rocky Mountain MS Brain Bank, Department of Neurology, University of Colorado School of Medicine.; Departments of Neurology and Neuroscience, Yale School of Medicine, Boyer Center for Molecular Medicine, 295 Congress Avenue, Ste 431-435, New Haven, CT, 06510.; Analytic Imaging Facility, Albert Einstein College of Medicine, 1300 Morris Park Avenue Bronx, New York 10461.; Graduate student. Department of Biochemistry, Albert Einstein College of Medicine, 1300 Morris Park Avenue Bronx, New York 10461, USA; UCLA Brain Bank. VA Healthcare System, 127A 11301 Wilshire Blvd., Los Angeles, CA 90073.

**Keywords:** Multiple Sclerosis, TMEM106B, myelin oligodendrocyte glycoprotein (MOG)-induced EAE, demyelination, lipids

## Abstract

During inflammatory, demyelinating diseases such as multiple sclerosis (MS) axonal damage is prevalent early in the disease course. Axonal damage includes swellings, defects in transport, and failure to clear damaged intracellular proteins, all of which affect recovery and compromise the integrity of neurons and remyelination. Autophagy and the clearance of damaged cell components by the proteasome are important for the maintenance of normal cellular turnover; and the restoration of cellular homeostasis. The gradual accumulation of insoluble proteins in the brain is known to impair recovery from several neurodegenerative diseases. In this study, we used mass spectrometry to identify insoluble proteins within high-speed, mercaptoethanol/sarcosyl-insoluble pellets from purified white matter plaques isolated from the brains of individuals with MS. We determined that insoluble transmembrane protein106B (TMEM106B), expressed in neurons and glia and normally lysosomal-associated, is increased in MS plaques relative to normal-appearing white matter from individuals with Alzheimer’s disease and non-neurologic controls. We found that decreased TMEM106B protein in mice results in significant axonal damage and lipid droplet accumulation in the spinal cord following chronic myelin oligodendrocyte glycoprotein-induced experimental autoimmune encephalomyelitis. When TMEM106B^t/t^ mice were treated with cuprizone to experimentally induce demyelination, a significant increase in lipid deposition was observed in the corpus callosum of TMEM106B^t/t^ mice post-cuprizone withdrawal. Our study shows that the brain and spinal cord from challenged TMEM106B^t/t^ mice accumulate OilRedO+/Perilipin2+ lipid droplets. We postulate that increased insolubility of TMEM106B in MS plaques limits debris clearance by the lysosome which over time contributes to failed remyelination and axonal defects.

**Abbreviated Abstract:** Transmembrane protein106B (TMEM106B), a lysosome-associated protein, is significantly less soluble in multiple sclerosis plaques than in white matter controls. Decreased TMEM106B produces significant axonal damage and lipid accumulation in mouse models of demyelinating diseases. TMEM106B insolubility and likely loss of function may limit lysosome transport and contribute to CNS pathology.

## Introduction

Multiple Sclerosis (MS) is a debilitating neurological disease with most individuals diagnosed between the ages of 20 and 50.^1,2^ In the United States there are an estimated one million individuals living with MS. During inflammatory attacks immune cells cross the protected blood-brain barrier and invade the nervous system, resulting in demyelination and axonal damage in brain and spinal cord. When prolonged by inefficient clearance of cellular damage and myelin debris, the repair processes are delayed. Over time, the progressive nature of MS results in a chronic disease course with permanent motor damage, demyelination, axonal damage, neuronal loss, OilRedO+ lipid inclusions and oligodendrocyte cell death. Thus, a major obstacle for individuals with MS is the maintenance of axonal function, and the ability to remyelinate following bouts of demyelination. We have identified increased levels of Transmembrane Protein 106B (TMEM106B) protein in sarcosyl-insoluble protein precipitates in white matter plaques from individuals with relapsing remitting multiple sclerosis (RRMS) relative to the white matter region of cortical normal-appearing sub-cortical white matter from non-neurologic controls. The increased insoluble TMEM106B led us to examine the consequence of reducing TMEM106B in mouse models of demyelination to determine whether the loss would impact spinal motor neurons during experimental autoimmune encephalomyelitis (EAE), a mouse model sharing pathological sequelae with MS, and impact remyelination in the cuprizone model of demyelination/remyelination.

TMEM106B is a 274 amino acid glycoprotein expressed in multiple cell types in the CNS including the lysosomal membrane of oligodendrocytes,^3^ microglia, astrocytes and neurons.^4–8^ The glycosylated carboxy-terminus faces the lysosomal lumen, and the 96-amino-acid amino-terminus is within the cytosol.^9^ The first 42 amino acids of TMEM106B are a region of high disorder permitting interaction with other proteins^9^. TMEM106B regulates lysosomal trafficking and lysosomal acidification.^9^ TMEM106B co-localizes with LAMP1 on lysosomes and binds progranulin and the microtubule-associated protein MAP6.^10^ The precise molecular function of TMEM106B is not known. Over-expression and deletion of TMEM106B *in vitro* result in varying pathologic consequences. *In vivo*,TMEM106B has opposing effects in mouse models of lysosomal diseases where it was found that TMEM106B differentially modulates the progression of the lysosomal storage diseases Gaucher disease and neuronal ceroid lipofuscinosis.^11^ Mice with deletion of TMEM106B and point mutations demonstrate functional defects in lysosomes, axonal damage, neurodegeneration, and hypomyelination. ^12–14^ With limited data on the role of TMEM106B *in vivo*, this study examined the function of TMEM106B in the CNS using TMEM106B(trap/trap; TMEM106B^t/t^) knockin mice, established by a gene trap insertion of lacz between exons 3 and 4 generating a hypomorphic knockin strain with reduced levels of total TMEM106B protein expressed.^11^. Our study shows that TMEM106B is significantly elevated in the insoluble fraction isolated from white matter plaques of individuals with relapsing remitting MS (RRMS). Further, when TMEM106B^t/t^ mice are sensitized with myelin-oligodendrocyte glycoprotein_35-55_ (MOG_35-55_) peptide lumbar spinal cords show increased lipid droplet accumulation, axonal damage and myelin loss relative to wildtype (WT) mice.

## Material and Methods

### Isolation of insoluble proteins from white matter

White matter lesions and normal appearing white matter were dissected from cortex and β-mercaptoethanol-, sarkosyl-insoluble proteins were purified using the isolation procedure from,^15^ followed by sonication, tryptic enzymatic digestion, before nanoLC-ms/ms analysis of the peptides using an Orbitrap Fusion Lumos mass spectrometer (Thermo Fisher Scientific). Sub-cortical white matter lesions from five individuals with relapsing remitting multiple sclerosis (RRMS) and normal-appearing sub-cortical white matter from non-neurologically impaired individuals were compared.

### Protein extraction and sample preparation

To analyze the proteome, the insoluble proteins in the pellet were processed for protein digestion using S-Trap filters (Protifi), according to manufacturer’s instructions. Briefly, the pellet was mixed with 5% SDS, followed by incubation with 5 mM DTT for 1 h, and then 20 mM iodoacetamide for 30 min in the dark, in order to reduce and alkylate proteins. Phosphoric acid was added to the samples at a final concentration of 1.2%. Samples were diluted in six volumes of binding buffer (90 % methanol and 10 mM ammonium bicarbonate, pH 8.0). After gently mixing, the protein solution was loaded to an S-Trap filter and spun at 500 g for 30 sec. The samples were washed twice with binding buffer. Finally, 1 μg of sequencing grade trypsin (Promega), diluted in 50 mM ammonium bicarbonate, was added into the S-trap filter and samples were digested overnight at 37°C. Peptides were eluted in three steps: (i) 40 μl of 50 mM ammonium bicarbonate, (ii) 40 μl of 0.1% trifluoroacetic acid (TFA) and (iii) 40 μl of 60% acetonitrile and 0.1% TFA. The peptide solution was pooled, spun at 1,000 g for 30 sec and dried in a vacuum centrifuge.

### Sample desalting

Prior to mass spectrometry analysis, samples were desalted using a 96-well filter plate (Orochem) packed with 1 mg of Oasis HLB C-18 resin (Waters). Briefly, the samples were resuspended in 100 μl of 0.1% TFA and loaded onto the HLB resin, which was previously equilibrated using 100 μl of the same buffer. After washing with 100 μl of 0.1% TFA, the samples were eluted with a buffer containing 70 μl of 60% acetonitrile and 0.1% TFA and then dried in a vacuum centrifuge.

### LC-MS/MS Acquisition

Samples were resuspended in 10 μl of 0.1% TFA and loaded onto a Dionex RSLC Ultimate 300 (Thermo Scientific), coupled online with an Orbitrap Fusion Lumos (Thermo Scientific). Chromatographic separation was performed with a two-column system, consisting of a C-18 trap cartridge (300 μm ID, 5mm length) and a picofrit analytical column (75 μm ID, 25 cm length) packed in-house with reversed-phase Repro-Sil Pur C18-AQ 3 μm resin. To analyze the proteome, peptides were separated using a 120 min gradient from 4-30% buffer B (buffer A: 0.1% formic acid, buffer B: 80% acetonitrile + 0.1% formic acid) at a flow rate of 300 nl/min. The mass spectrometer was set to acquire spectra in a data-dependent acquisition (DDA) mode. Briefly, the full MS scan was set to 300-1200 m/z in the orbitrap with a resolution of 120,000 (at 200 m/z) and an AGC target of 5×10e5. MS/MS was performed in the ion trap using the top speed mode (2 sec), and AGC target of 1×10e4 and an HCD collision energy of 30.

### Proteomics data analysis

Proteome raw files were searched using Proteome Discoverer software (v2.4, Thermo Scientific) using SEQUEST search engine and the Swissprot human database (updated February 2020). The search for total proteome included variable modification of N-terminal acetylation and fixed modification of carbamidomethyl cysteine. Trypsin was specified as the digestive enzyme with two missed cleavage allowed. Mass tolerance was set to 10 ppm for precursor ions and 0.02 Da for product ions. Peptide and protein false discovery rate was set to 1%. Prior to statistical analysis, proteins were log2 transformed, normalized by the average value of each sample, and missing values were imputed using a normal distribution 2 standard deviations lower than the mean. Statistical regulation was assessed using heteroscedastic T-test (if p-value < 0.05).

### Stains and Antibodies

The Brain Stain Imaging kit (B34650; Molecular Probes) was used to stain myelin in frozen brain sections. The kit also stains nuclei, and neurons (Nissl). Antibody SMI32 (Millipore; 1:10,000) recognizes non-phosphorylated neurofilament protein in axons. SMI99 (Millipore; 1:1000) recognizes myelin basic protein (MBP). Iba1 (WAKO, for IHC; 019-19741; 1:400) recognizes microglia/macrophage. An affinity-purified TEM106B antibody was purchased from Proteintech (rabbit polyclonal 290995-1-AP; 1:100). Perilipin 2 (PLIN2) antibody recognizes the N-terminal amino acids 1-29 (Progen GP40, 1: 400). APC/CC1 (OP80 Millipore; 1:20) recognizes mature oligodendrocytes. Carbonic anhydrase II (CAII; ab124687; Abcam; 1:250) recognizes oligodendrocytes. LAMP1 antibody (1D4B; 1:100) was purchased from Developmental Studies Hybridoma Bank, University of Iowa). LC3(A/B) (D3U4C) XP Rabbit monoclonal antibody was purchased from Cell Signaling (12741). All Alexa-fluorescent secondary antibodies were purchased from Fisher Scientific.

### Immunostaining and analysis of human brain sections

Fresh frozen MS and non-neurological control tissue (Rocky Mountain Brain Bank) was used for mass spectrometry and some samples were prepared for immunostaining by formalin fixation and paraffin-embedding. Addition formalin-fixed tissue was obtained from the UCLA Brain Bank and was paraffin-embedded, or embedded in O.C.T. Compound (4583; Tissue-Tek); 5-7 micron sections were prepared. Alzheimer brain tissue was obtained from the New York Brain Bank at Columbia University Irving Medical Center, and additional material was obtained from the Montefiore Brain Repository. Following staining of human brain sections, slides were scanned on a P250 slide scanner and full sections were evaluated and photographed in Caseviewer. At least 30 random fields of white matter and 20 of gray matter were examined for all cases and representative fields were photographed at x63 white matter, x40 gray matter. Figure legends detail the staining and antibody.

### Mice

TMEM106B^t/t^ mice on a Bl6 background were obtained from Dr. Stephen Stritmatter at Yale University^16^ and were used to examine two mouse models of CNS damage, the cuprizone model of demyelination/remyelination and myelin oligodendrocyte glycoprotein (MOG)-induced experimental autoimmune encephalomyelitis (EAE). The strain TMEM106B (trap/trap; TMEM106B^t/t^), was established by a gene trap insertion of lacz between exons 3 and 4 generating a hypomorphic knockin strain herein referred to as TMEM106B^t/t^ Low amounts of the full-length protein is detected by Western blot analysis^11^. The genotypes were confirmed by Transnetyx using duplexed PCR assays for both WT and knockout (KO) alleles.

P017: 5’ GGGATCTCATGCTGGAGTTCTTCG 3’

P019: 5’ TTCTCTCCATGTGCTGCATTATGAGC 3’

P020: 5’ ACGTGCTTCTCTCATCTAGAGTTTTCC 3’

The TMEM106B^t/t^ mouse model differs from models with a complete deletion of TMEM106B.^9,12–14^ Unlike our strain, a complete knockout of TMEM106B showed motor deficits at 5.5 months of age, while other groups reported oligodendrocyte deficits using the cuprizone model.^13,14^

#### Ethical Statement for Animals, housing and husbandtry, care and monitoring

All animals will be bred, maintained, and treated in the Barrier Facility under the guidance of the Albert Einstein College of Medicine (AECOM) Veterinary staff and Dr. Shafit-Zagardo approved protocol number 00001158. All procedures are in complete compliance with the AECOM Institutional Review Board and with the NIH Guide for the Care of Laboratory Animals. Mice were continually monitored for any distress including weight loss, changes in excretion, or lethargy. Wherever possible such as scoring mice the testers were blinded to the genotypes of the groups. Experiments are repeated at least twice to ensure consistent findings. Mice undergoing EAE are monitored daily and provided with standard chow pellets and DietGel Recovery (clearh2.0com) at the cage bottom if they show hindlimb weakness or paralysis. Mice are sacrificed if they score 5 moribund. Each animal acquires significant value that is protected by good animal care and by ensuring the animal’s comfort. The approved euthanasia protocol employed was outlined by the Panel on Euthanasia of the American Veterinary Medical Association and IACUC offices. To sacrifice moribund animals mice receive isofluorane in a sealed chamber, and perfused with 4% paraformaldehyde. Both males and females were included in the studies, unless otherwise specified.

### MOG-induced EAE

C57Bl6J and TMEM106B^t/t^ mice were immunized with myelin oligodendrocyte glycoprotein (MOG)_35-55_ peptide emulsified in complete Freund’s adjuvant and injected with pertussis toxin on day 0 and day 2^17^. Mice were monitored daily for clinical symptoms and were scored as follows: 0 = no clinical symptoms, 1 = flaccid tail, 2 = flaccid tail and hind limb weakness, 3 = hind limb paralysis, 4 = hind limb paralysis and forelimb weakness, 5 = moribund. Mice that did not present with clinical symptoms were excluded from analysis.

### Cuprizone Treatment

Mice were fed 0.2% *(w/w)* cuprizone (Sigma; St. Louis, MO) in powdered chow *ad libitum* for 5 weeks, followed by regular chow pellets. Mice were sacrificed at 5-weeks to confirm demyelination, and during the recovery phase, 2-, or 3-weeks after the removal of cuprizone from the diet. For immunohistochemical analysis, brains were removed following cardiac perfusion with 4% paraformaldehyde, and paraffin-fixed and frozen sections were prepared. Coronal sections of brain (10 μm) were cut on a cryostat and the midline of the corpus callosum was evaluated at the region of the fornix corresponding to sections 230–260 of Sidman’s mouse atlas ^18^.

### Oil Red O staining

Frozen sections were incubated in distilled water for 1 min followed by a 2-min incubation in 100% propylene glycol (Polyscientific; Bayshore, New York) and immediately transferred to Oil Red O stain (Polyscientific) for 48 hours at room temperature. Sections incubated for 1 min in 85% propylene glycol, rinsed in double-distilled deionized (dd)H_2_O for 1 minute, and mounted with glycerin jelly mounting medium (Polyscientific). Mounted cross sections were scanned on a 3D P250 High Capacity Slide Scanner. Slides were imaged with a 4-megapixel-color CMOS camera with a 20x objective and then stitched together. Images were analyzed using Case Viewer software.

### Histologic grading

Histological sections were graded on a scale of 0–4 as previously described^19^ For Iba1^+^ staining, cross-sectional spinal cords or brains were scored on a 0–4 inflammatory scale where a score of 0 is the equivalent pathology observed in a naïve mouse, 1 = mild inflammation, 2 = moderate, 3 = severe inflammation, and 4 = very severe inflammation involving 50% or more of the tissue. For relative changes in myelination and to assess decreased MBP staining, the scores were assigned as follows: 0 = MBP immunoreactivity observed in naïve mice, 1 = mild demyelination, 2 = moderate demyelination, 3 = severe demyelination, and 4 = very severe involving >50% of white matter. For relative axonal damage, we either counted all SMI32+ axons in the white matter or quantified the SMI32^+^ axonal swellings (>3 μm) in the left and the right ventral region of the white matter of mouse spinal cords using multiple 20× fields. 0 = 0 SMI32^+^ axonal swellings as observed in naïve mice, 1 = ≤ 10 SMI32^+^ swellings, 2 = 10–20 SMI32^+^ swellings, 3 = 20–50 SMI32^+^, and 4 = ≥50 SMI32^+^ swellings. Slides were blinded and at least three sections of lumbar spinal cord for each animal were assessed by two individuals; the number of animals used for each experiment is indicated in the figure legends. Mann-Whitney *β*-test was used to evaluate statistical significance.

### Statistical Analysis

All values are presented as mean or Median ± standard error of the mean (SEM). Statistical analyses were performed in GraphPad Prism Software. Significance was determined using a two-tailed unpaired student’s t-test for parametric analyses and Mann-Whitney for nonparametric analyses. Multiple group analysis was perfromed using One-way ANOVA.

### Data Availability

The authors confirm that the data supporting the findings of this study are available within the article and its supplemental material.

## Results

### TMEM106B is elevated in insoluble protein pellets isolated from white matter plaques of individuals with RRMS

Mass spectrometry was used to determine the composition of β-mercaptoethanol/sarkosyl-insoluble proteins within sub-cortical white matter of MS plaques relative to those in normal-appearing white matter. We hypothesized that these insoluble proteins are possibly sequestered or unavailable for normal function. While the mass spectrometry studies are still ongoing, we examined several of the proteins in more detail, including TMEM106B, which we determined to be significantly enriched in the pellet from RRMS plaques (red dot, Fig. 1A); p=0.0071, Mann-Whitney U test). The log2 TMEM106B peptides represented in insoluble pellets from non-neurological controls (n=5) and RRMS (n=5) samples were examined from one experiment in which all 10 samples were processed and run at the same time. Both male and female samples showed no difference in scores within the group. Subsequent groups containing additional male and female samples confirmed our findings. The unique peptides identified for TMEM106B are shown in Fig.1 B,C. A Bean plot displays the differences between the non-neurological white matter controls (n=4; one outlier) and those from RRMS plaques (n=5; Fig. 1D). The data are presented as median ± interquartile range (1^sl^ quartile, 3^rd^ quartile). The median value for the non-neurologic controls was log_2_ 25.79 (25.04, 26.46), and the log2 RRMS was 30.26 (29.40, 30.90). TMEM106B presence was confirmed and quantified using four peptides confidently identified by mass spectrometry (False Discovery Rate <0.01).

**Figure 1:**
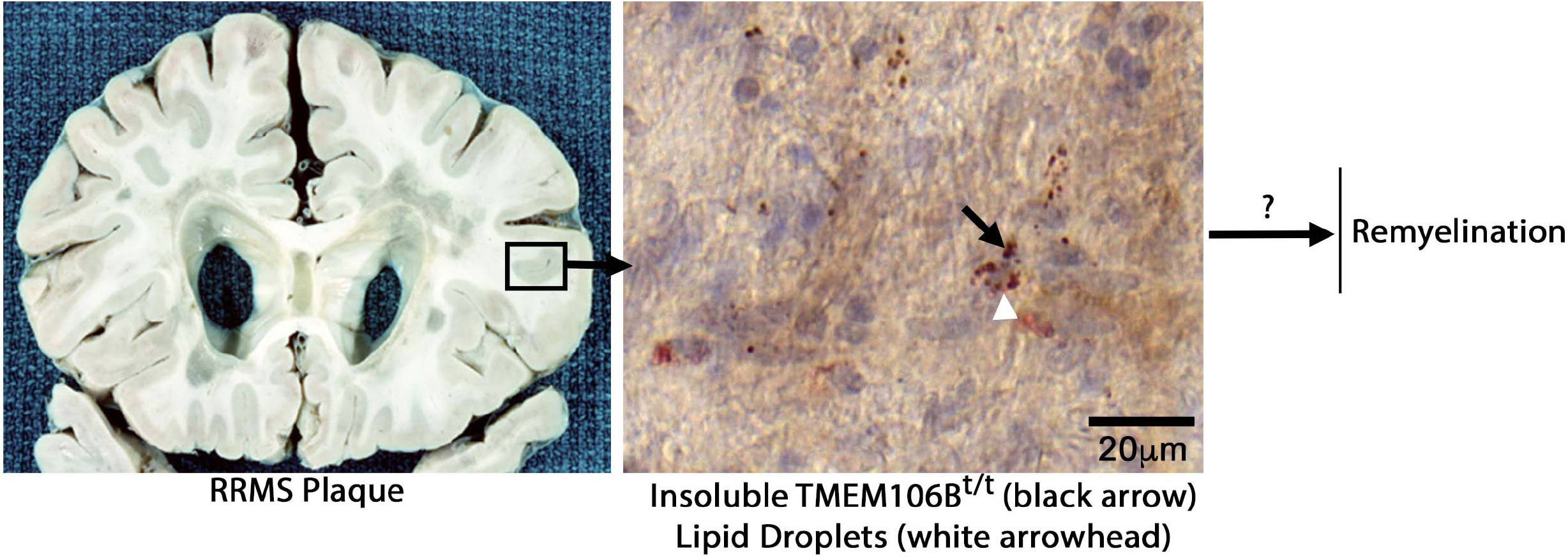
TMEM106B is elevated in protein pellets isolated from white matter plaques of individuals with RRMS. Identification and quantification of TMEM106B via mass spectrometry-based proteomics. **(A)** Volcano plot displaying log2 fold change and -log2 t-test p-value for the comparison of RRMS vs Control samples. TMEM106B is highlighted in red. The distribution of the data points shows upregulated and downregulated proteins, including several non-significant (significance threshold set at 4.32 = -log2(0.05)). The mean log2 of TMEM106B peptides identified in the sarcosyl-insoluble protein pellets from plaques of individuals with RRMS relative to non-neurologic controls following nanoLC-ms/ms analysis of peptides on an Orbitrap Fusion Lumos mass spectrometer (Thermo Fisher Scientific). The mean age of the non-neurologic control group was 63.2 ±3.0, the RRMS was 64.8 ±1.3 years of age. The mean age of each group, and the post-mortem intervals were not significantly different. Mann Whitney U-test, 24, p=0.5476. **(B)** Coverage of the protein TMEM106B based on confidently identified peptides. The first two peptides are the same region of the protein, but they differ for one missed cleavage of the tryptic digestion. **(C)** Annotated MS/MS spectra for each of the identified peptides. Fragment ion signals highlighted in red represent the b-series fragments (N-terminus), while those in blue are the y-series fragments (C-terminus). **(D)** Quantification of the protein TMEM106B using the average signal intensity of the four identified peptides. From left to right: protein abundance for each of the replicates as bar plot, raw values as table, bean plot representation of the log2 transformed values, outlier analysis of the data distribution. The upper bound of the Ctrl distribution is smaller than the first replicate value, therefore we categorized it as an outlier. Without the outlier, the Mann Whitney test is 0.0071; including the outlier the p value was 0.035.

### White matter plaques of individuals with RRMS show increased TMEM106B+ puncta relative to white matter controls

TMEM106B immunoreactivity was examined using paraffin-embedded brain sections containing MS plaques, adjacent white matter, and gray matter from individuals with RRMS. Consistent with the presence of TMEM106B protein in sarcosyl-insoluble pellets, we observed numerous TMEM106B+ puncta in plaques and adjacent white matter in RRMS white matter. Plaques from an individual with SPMS also showed TMEM106B+ puncta (Fig. 2 and Supplemental Table 1). When compared to white matter from individuals with Alzheimer’s disease (n=6), and non-neurologic controls (n=10), the number of cells with TMEM106B+ puncta were increased/x63 fields in RRMS plaques (8.65±3.64; median±SEM; n=6), compared to AD white matter cases showing fewer than 3 cells/x6x white matter field (1.35±0.326; median±SEM; n=6), in 30 random fields/individual examined. Examiniation of sub-cortical white matter from individuals who had no neurologic involvement at the time of death, showed lower numbers of TMEM106B+ cells/x63 microscopic fields (1.94±0.814; median±SEM; n=10), and fewer TMEM106B+ puncta/cell than MS samples. The TMEM106B+ puncta/cell was similar to the number of TMEM106B+ puncta/cell observed for Alzheimer disease white matter. In general, the number of puncta/cell was increased in the MS plaques with ~4-8 puncta/cell (5.9±0.81) compared to Alzheimer disease white matter cells containing ~2-3 puncta/cell.

**Figure 2.**
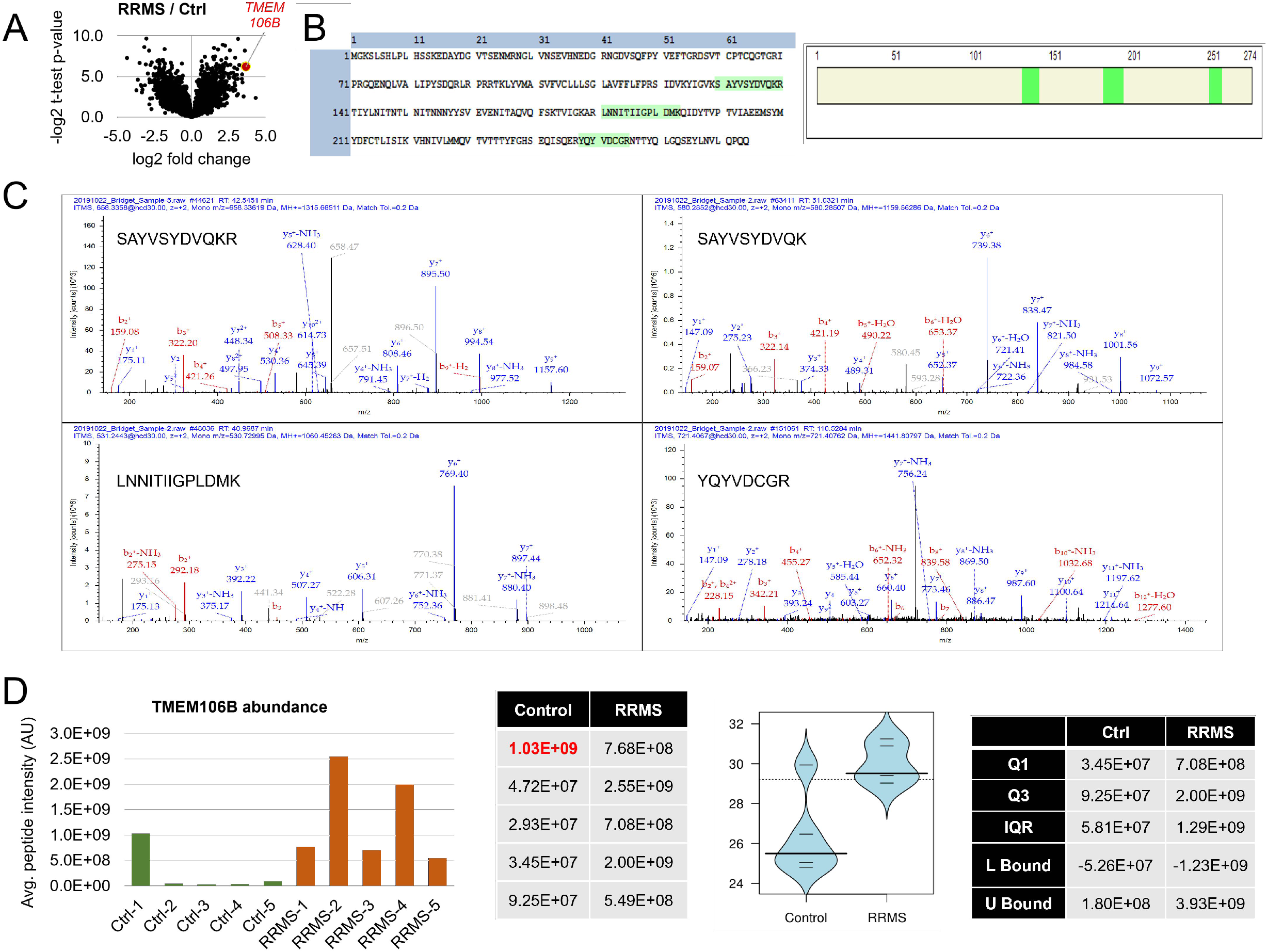
Increased TMEM106B in RRMS plaques. **(A)** Sub-cortical white matter sections from RRMS and SPMS brains have abundant TMEM106B+ puncta. **(B)** Statistical analysis of TMEM106B+ cells/x63 field, 30 fields was performed using the Kruskal-Wallis test followed by post-hoc pairwise comparisons using Mann-Whitney U-tests. The Kruskal-Wallis test statistic =16.74; p=0.0008. Multiple comparisons adjusted p values RRMS vs AD=0.0188. RRMS vs NNC=0.0088. Gray dots in the non-neurological control sample represent the septic shock and pneumonia samples. **(C)** TMEM106B+ neurons in gray matter. Over 30 random white matter fields were examined at x63, and 20 random x40 gray matter fields. Visualization was by diaminobenzidine (DAB). Arrowhead denotes the axon and arrow the apical dendrite.

Examination of cortical gray matter of individuals with MS, Alzheimer disease, and non-neurological controls showed TMEM106B+ punctate staining in neuronal cell bodies, dendrites, and axons (Fig. 2B). Pronounced punctate staining was apparent in neurons and processes of individuals with RRMS, and often we observed more diffuse TMEM106B+ staining in neuronal cell bodies, dendrites, and axons of non-neurological controls.

While the mass spectrometry data show TMEM106B is enriched in insoluble pellets in RRMS, and immunostaining shows increased numbers of TMEM106B+ cells/field, post-mortem MS material does not allow for the type of analysis mouse models permit. Therefore, we examined TMEM106B^t/t^ mice in two models of nervous system injury, MOG-induced EAE and cuprizone-induced demyelination and remyelination. Our prediction was that a reduction in TMEM106B levels would result in more axonal damage in TMEM106B^t/t^ mice during chronic EAE, with OilRedO+ deposition indicating that myelin and cellular debris are not efficiently cleared by the lysosome following injury in both model systems.

### Naïve WT and TMEM106B^t/t^ mice exhibit similar myelination and CNS architecture at the corpus callosum and spinal cord

Prior to performing studies, we analyzed the brains and spinal cords of naïve C57B6J and TMEM106B^t/t^ mice for any deficits in 8-10 week young adult mice and at 5.5 months of age. We determined that TMEM106B^t/t^ mice have normal cellular architecture, normal appearing gray matter with equivalent motor neurons in spinal cord. A myelin stain (Brain Stain Imaging kit (B34650) of the corpus callosum and the spinal cord is shown in Supplemental Figure 1. On outward inspection, the mice had no gait abnormalities or weakness on a hanging test.

### TMEM106B^t/t^ mice fail to efficiently clear myelin debris, and have significant axonal damage and demyelination during chronic EAE

MOG-induced EAE is a pertinent model to validate and characterize proteins in the context of inflammatory demyelination. To examine the consequence of reduced TMEM106B during EAE, TMEM106B^t/t^ and WT mice were sensitized with MOG35-55 peptide and monitored over the course of EAE. There was no difference in the day of onset of clinical scores or indices (CI) between the WT (12.6±0.75 days) and the TMEM106B^t/t^ mice (11.0±1.2 days) when individual groups of male and female mice (n= ~10/group) were challenged. No significant differences in the clinical indices of WT and TMEM106B^t/t^ were observed. The same clinical course was observed for both groups of mice during acute and chronic disease (Fig. 3). Mice at approximately day 42 post-MOG injection were sacrificed; the mean clinical score per group was 1.17±0.17 [WT] and 1.63±0.47 [TMEM106B^t/t^]. When lumbar spinal cord sections were examined histologically, there was no significant difference in the overall number of lesions assessed by H&E, the extent of gliosis as assessed for GFAP+ astrocytes, or Iba1+ activated microglia (not shown).

**Figure 3:**
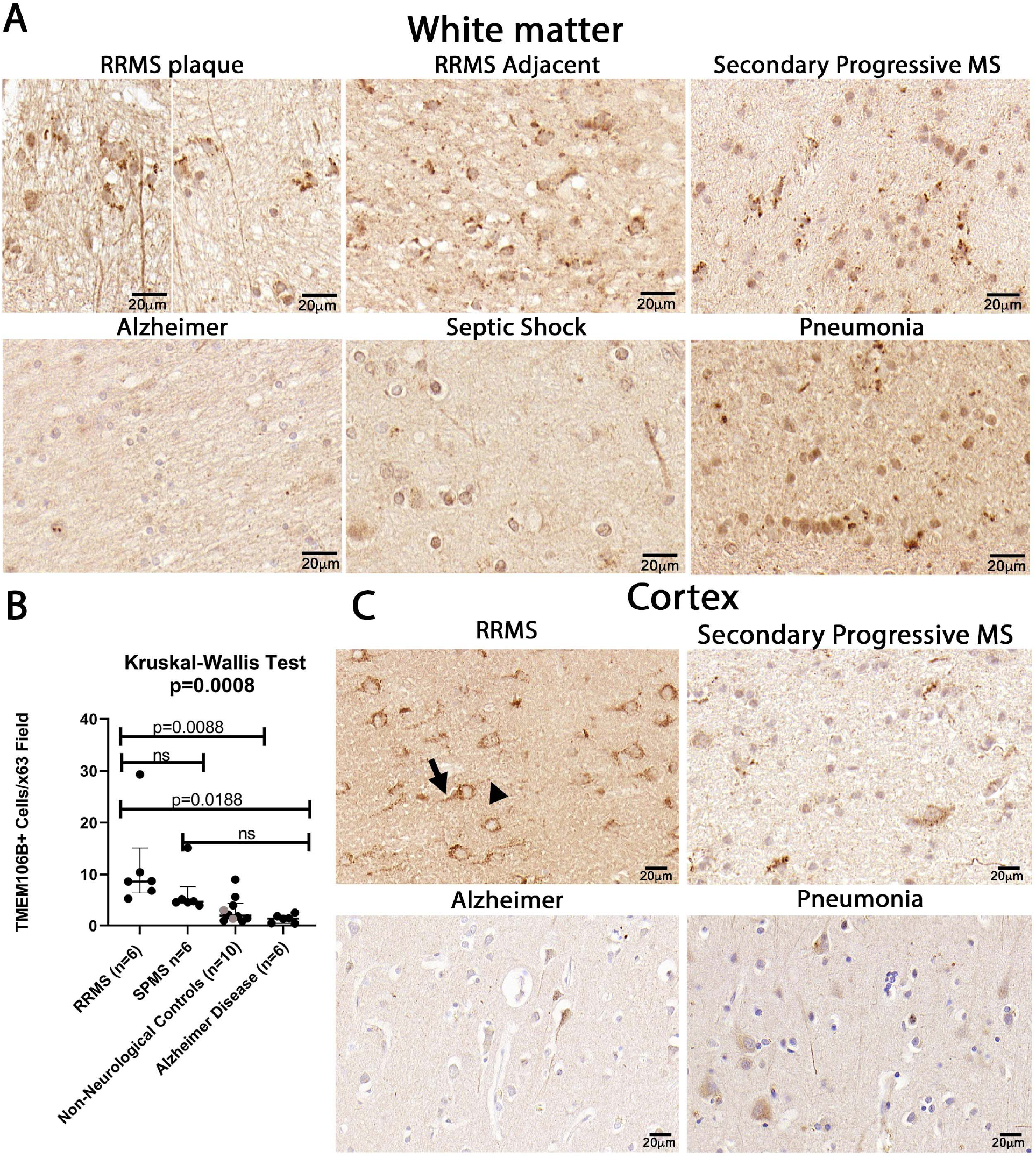
TMEM106B^t/t^ spinal cords show increased OilRedO+ lipid deposition and greater loss of myelin than sex- and age-matched control mice with the similar clinical scores throughout the clinical course. No differences in male and female mice were observed between the two groups of mice. OilRedO staining of naive female WT **(A)** and TMEM106B^t/t^ **(B)** spinal cords, and chronic WT **(C)** and TMEM106B^t/t^ **(D)** spinal cords at ~ 42 days post-MOG injection. Images from chronic EAE are seen in **(C-L). (F,H)** shows OilRedO staining at the central canal. Increased OilRedO deposition was observed in neurons from TMEM106B^t/t^ lumbar spinal cords **(I,J)** relative to WT spinal cord **(G)**. A myelin stain (Brain Stain Imaging kit B34650; Molecular Probes) shows reduced myelin TMEM106B^t/t^ spinal cord (**L**, green) relative to control (**K**). Nuclei are stained blue (dapi), neurons are stained magenta (Nissl) and lipids (green).

Cross sections of spinal cords were examined for OilRedO+ deposition of neutral lipid droplets, myelin loss using a myelin stain (Brain Stain Imaging kit B34650; Molecular Probes) and myelin basic protein (MBP) immunoreactivity, and axonal damage was assessed by SMI32+ swellings. To assess the extent of efficient clearance of neutral lipid debris in the two groups of mice, we stained frozen spinal cord sections with OilRedO. Naïve WT and TMEM106B^t/t^ mice have no OilRedO deposition in the spinal cord at 5.5 months of age (Fig. 3A,B); all MOG-sensitized mice were less than 4.5 months of age at the time of sacrifice. While there is minimal to no OilRedO deposition in all the WT spinal cords examined during chronic EAE, there were OilRedO+ lesions remaining in TMEM106B^t/t^ spinal cords both in white and gray matter (Fig. 3C-D). The ventral region of the spinal cord with OilRedO+ staining in Fig. 3D (arrowhead) is magnified in Fig. 3E. The mice were matched for similar clinical course throughout EAE, final chronic clinical score, and for sex. We detected OilRedO+ staining at the central canal in both WT and TMEM106B^t/t^ mice (Fig. 3F,H). The presence of OilRedO+ lipid deposits within spinal neurons of TMEM106B^t/t^ mice was apparent during chronic EAE (Fig. 3I,J). This is in contrast to the lack of lipid deposits in spinal neurons in the WT mice (Fig. 3G).

A myelin (green), Nissl (magenta), DAPI (blue) stain showed the extent of demyelinated regions in frozen crosssections of WT and TMEM106B^t/t^ spinal cords (Fig. 3K,L). To further quantify changes in myelin and axonal damage in the mice, we stained paraffin-embedded spinal cord sections with an antibody to myelin basic protein (MBP; SMI99), and assessed axonal damage using an antibody to non-phosphorylated neurofilament protein (SMI32). There was a significant difference in MBP staining (green) in the spinal cords of TMEM106B^t/t^ mice when myelin loss was scored on a 1-4 scale (Fig. 4). The demyelinated ventral region of TMEM106B^t/t^ lumbar spinal cord show a significant increase in the number of SMI32+ swelling (magenta) in lumbar spinal cord (Fig. 4A,B). The SMI32+ swellings are observed on both naked axons devoid of myelin, and axons still ensheathed by myelin (arrow), indicative of ongoing axonal injury. The total number of SMI32+ axonal swellings within the entire cross-section of the lumbar spinal cord was significantly higher in TMEM106B^t/t^ mice relative to WT (Fig. 4C; p=0.030). The relative degree of demyelination as assessed by SMI99+ staining in the TMEM106B^t/t^ mice relative to WT mice was also significant (p=0.028; Fig. 4D). No significant differences in Iba1+ glia was observed between WT and TMEM106B^t/t^ mice using a relative Iba1 inflammatory score (0-4; Fig. 4E-G; Mann-Whitney-U test 10.50, p=0.303), or the mean fluorescent intensity /area was measure for each lumbar spinal cord region; Kruskal Wallis statistic 8.467; p=0.1323 (data not shown).

**Figure 4:**
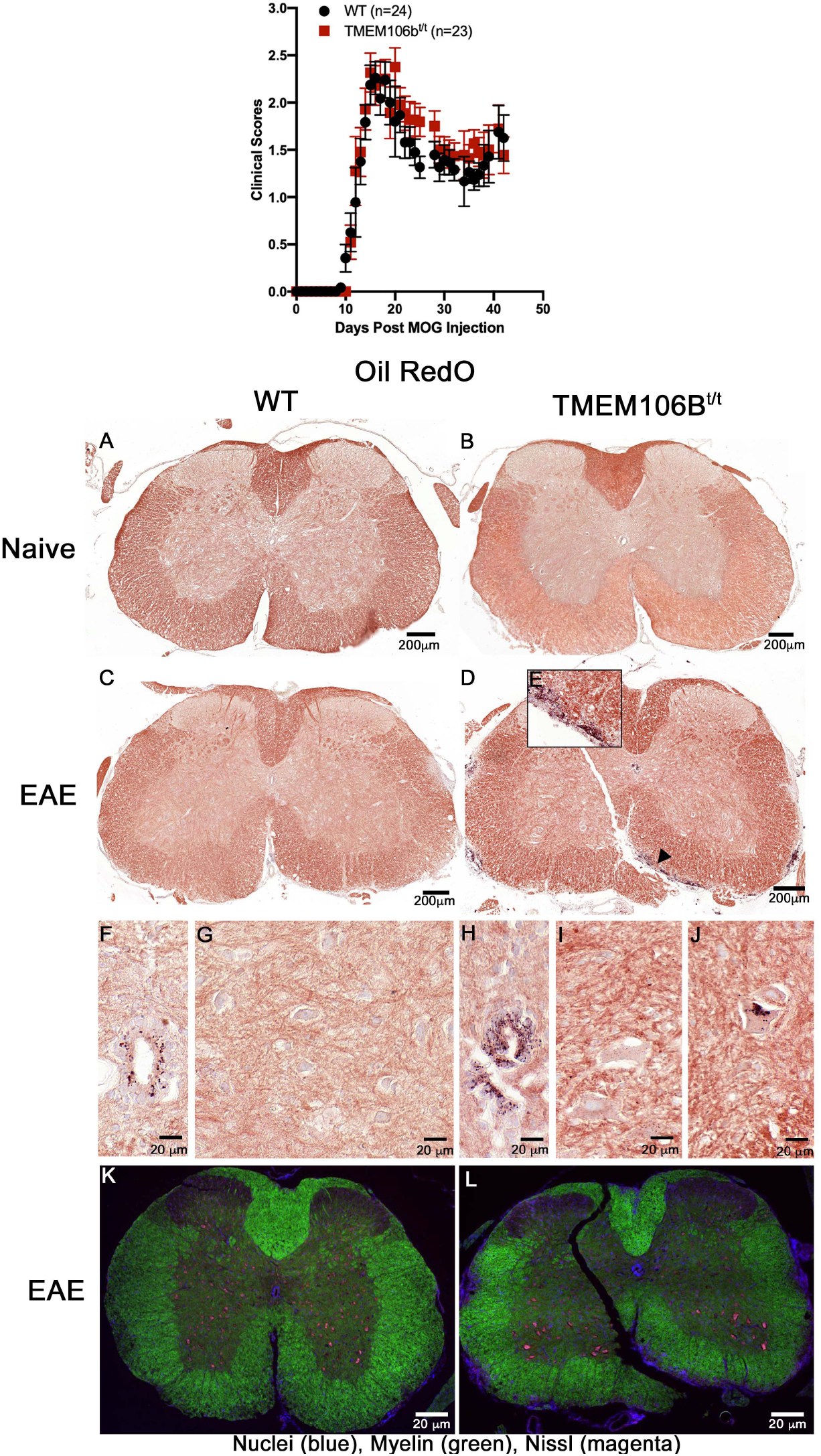
TMEM106B^t/t^ spinal cords show more axonal damage and demyelination relative to female control mice. **(A)** WT (n=5) and **(B)** TMEM106B^t/t^ (n=6) spinal cord sections were staining with dapi (blue), MBP (green), and SMI32 (magenta) during chronic EAE. Shown is the ventral region of lumbar spinal cord. **(C)** SMI32+ swelling/cord were counted and quantified. Mann-Whitney U-test 3, p=0.0303. **(D)** Relative SMI99 (MBP) was scored on a 0-4 scale. Mann-Whitney U-test 3.5, p=0.0281. **(E)** Iba1inflamatory score showed no differnces in the two groups during chronic EAE. Statistical anlaysis was performed using GraphPadPrism Software. Mann-Whitney U-test. Iba1 Mann-Whitney U-test 10.5, p=0.303. Represntative Iba1 immunofluorescent staining (orange) of WT **(F)** and TMEM106B^t/t^ spinal cord **(G).**

As a result of the increased OilRedO+ lipid deposits observed at the central canal and spinal cord of TMEM106B^t/t^ mice during chronic EAE, we incubated additional spinal cord sections with an antibody to perilipin2 (PLIN2), a resident lipid droplet-protein. The central canal of both WT and TMEM106B^t/t^ mice were positive for PLIN2 (Fig. 5A,B; green); no primary antibody was used as a control to stain WT spinal cord (Fig. 5C). Relative to WT spinal cord (D,E), PLIN2+ cells in gray and white matter of TMEM106B^t/t^ spinal cord were prevalent (F-H). The *F shows an intensely positive PLIN2 cell from an adjacent gray matter section. The clinical scores, sex, and duration of chronic disease were the same in the WT and TMEM106B^t/t^ mice that were compared. A no primary control of TMEM106B^t/t^ white matter is shown in Fig. 5I.

**Figure 5:**
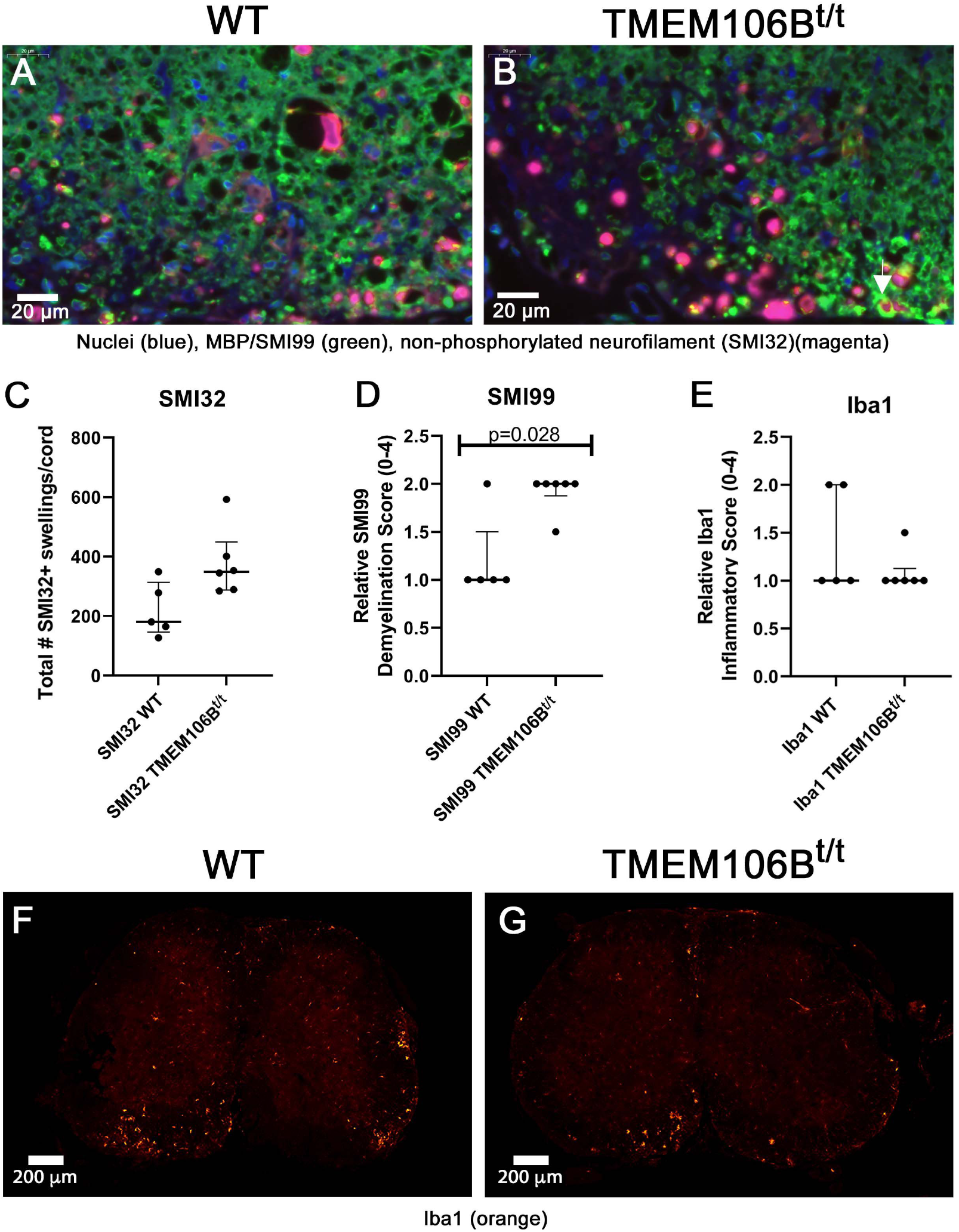
Perilipin2 (PLIN2) is expressed at the central canal and in spinal cord during chronic EAE. **(A)** Central canal of female WT and TMEM106B^t/t^ mice 45 days post-MOG injection. **(B)** WT neurons and glia in lumbar spinal cord (**D,E**) have less PLIN2 staining than TMEM106B^t/t^ (**F,G,H**). Insert **F(*)** shows an intensely positive PLIN2+ cell from an adjacent gray matter section; (n=3/group). No primary antibody control (**C,I**).

To further investigate the cause of the lipid droplet accumulation in the CNS of mice, we sensitized a small group of WT and TMEM106B^t/t^ mice with MOG peptide and examined the lipid droplet accumulation in spinal cord when the mice had clinical scores CI≥1, for 18 consecutive days (Fig. 6A, late acute EAE). Again, there was no significant difference in the clinical scores of the two groups of mice. Frozen spinal cord sections from WT mice all with a final CI=1 (n=3) and TMEM106B^t/t^ (n=5; CI=1-2) mice were incubated with OilRedO. Mice with CI=1 were paired based upon the number of days with the same final clinical score, and the number of days throughout the disease course with CI=2’s and 3’s, that would contribute to worsening of spinal cord damage. When compared to the OilRedO+ spinal cords during chronic EAE, there were fewer OilRedO+ cells at the central canal of WT and TMEM106B^t/t^ mice, and there was minimal OilRedO+ staining in the three WT spinal cords (Fig. 6). TMEM106B^t/t^ spinal cords having CI=1 and similar courses to the WT mice, showed OilRedO+ staining in neurons (Fig 6E, arrow). A spinal cord from a TMEM106B^t/t^ mouse with a final score of CI=2 had more OilRedO staining in the ventral and lateral spinal cord, and in neurons (Fig. 6F,G), similar to the spinal cords of TMEM106Bt^4^ mice during chronic EAE. PLIN2 staining in the spinal cord of the WT mice CI=1 showed staining in neurons, glia and at the central canal (Fig. 6H,I). The number of PLIN2+ lipid droplets in neurons in the TMEM106B^t/t^ did not significantly increase relative to WT (Fig. 6J,K), as some of the WT PLIN2+ neurons had equivalent numbers of PLIN2+ droplets; Kruskal-Wallis statistic 9.052, p=0.107. The PLIN2+ cells in spinal cord of the TMEM106B^t/t^ mouse with CI=2 paralleled the OilRedO+ staining observed in gray matter and ventral white matter regions (L,M). Our combined OilRedO and PLIN2 studies show that there is an accumulation of lipids in TMEM106B^t/t^ spinal cords during chronic EAE that can be seen during early recovery. Qualitatively, the higher the clinical scores reflective of greater CNS damage, the more OilRedO+PLIN2+ cells detected.

**Figure 6.**
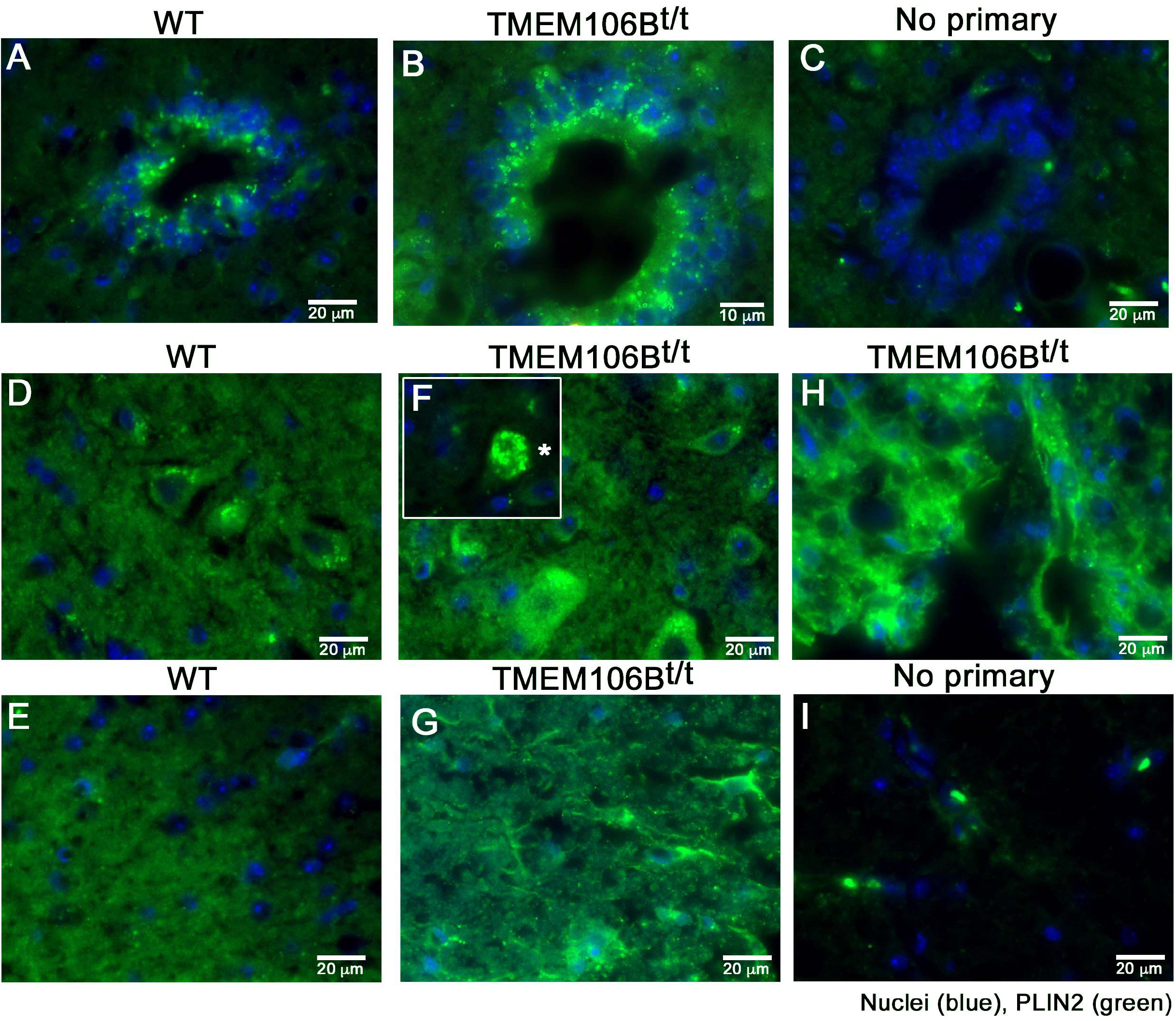
TMEM106B^t/t^ mice sensitized with MOG35-55 have OilRedO+ lipid deposits at the central canal and in spinal cord after consecutive clinical scores for 18 days. (**A**) No significant difference in the clinical scores of female WT (black) and TMEM106B^t/t^ (red) mice. (**B**) OilRedO+ staining of frozen spinal cord sections of WT mice CI=1 (n=3; **B,C**), and TMEM106B^t/t^ mice CI=1 (n=3; **D,E**) and CI=2 (n=1; **F,G**). (**H-O**) PLIN2 immunostaining (green) including axons.

### TMEM106B^t/t^ mice have significant OilRedO+ deposits in the corpus callosum at 2- and 3-weeks post-cuprizone withdrawal

10-week old WT and TMEM106B^t/t^ male and female mice were fed 0.2% cuprizone in powdered chow for 5-weeks, and groups of mice were sacrificed at 2- and 3-weeks post-cuprizone withdrawal. Based on our analyses during EAE, we predicted that the corpus callosum of TMEM106B^t/t^ mice would retain OilRedO+ deposition throughout recovery. TMEM106B^t/t^ mice had significantly more and larger OilRedO+ inclusions at the midline and lateral regions of the corpus callosum at 2-weeks post-cuprizone withdrawal when there was minimal deposition in the WT corpus callosum (Fig. 7C,D). Shown are the midline of WT and TMEM106B^t/t^ corpora callosa (n=3/group). Less than 10% of the area of WT corpora callosa contained OilRedO+ puncta when examined at x40 using CaseViewer. Images were divided into four quadrants, and small puncta with a diameter of <3 μm were observed in one to two quadrants of the WT corpora callosa. Examination of the lateral regions showed few OilRedO+ puncta of <2 μm. OilRedO+ puncta were detected throughout all four quadrants of the midline of the corpus callosum of TMEM106B^t/t^ mice, and the size of the puncta were larger. OilRedO+ puncta ranging from 3.8-8.9 μm were detected in all quadrants. In the lateral regions, large puncta from 3.8-8.6 μm were observed. By 3-weeks recovery, there were still large OilRedO+ inclusions at the midline of TMEM106B^t/t^ corpus callosum and the lateral regions. The WT mice had no visible OilRedO+ puncta at the midline and only one of three had small puncta in the lateral regions of the corpus callosum (Fig. 7E,F). The corpus callosum of control 5.5 month naïve WT and TMEM106B^t/t^ mice appeared normal with no OilRedO+ deposits (Fig. 7A,B). The myelin stain (Brain Stain Imaging kit) showed no significant difference in myelin in WT and TMEM106B^t/t^ corpora callosi at 2- and 3-weeks recovery (Fig. 7G-J). Examination of oligodendrocyte protein markers showed no significant difference in oligodendrocyte number as assessed by APC/CC1 immunofluorescent staining (Fig. 7K,M). Carbonic anhydrase (CAII), another oligodendrocyte marker for oligodendrocytes that participates in the myelination of small caliber axons ^20^ showed no difference in oligodendrocyte number between the two groups (Fig. 7K,M). SMI32+ staining of axons at both 2- and 3-weeks post-cuprizone withdrawal showed no significant differences in the number of axonal swellings (Fig. 7K,L). Iba1+ score was no different at 2 week recovery (K), and there was no difference in the extent of gliosis (data not shown). MBP mean fluorescent intensity was not different between the two groups at 3-weeks recovery (Fig. 7N). Since lipid droplets can co-localize with the autophagy marker LC3 ^21^, we measured the number of cells with LC3+ puncta as a ratio of the total number of cells in the corpus callosum per 60x field at 3-weeks post-cuprizone withdrawal. There was an increase in the percentage of cells containing LC3+ puncta in TMEM106B^t/t^ corpus callosum (22.7±1.1) relative to WT mice (11.8±2.0); p=0.0074, (Mean±SEM; n=3-4/ group; unpaired t-test). Additional sections yielded similar results, TMEM106B^t/t^ corpus callosum (21.0±1.1), WT corpus callosum (15.3±0.96); p=0.0107. (Mean±SEM; n=3 TMEM106B^t/t^, WT n=4; unpaired t-test). Due to the small number of samples, when a Mann-Whitney U-test was performed the p value was p=0.0571.

**Figure 7:**
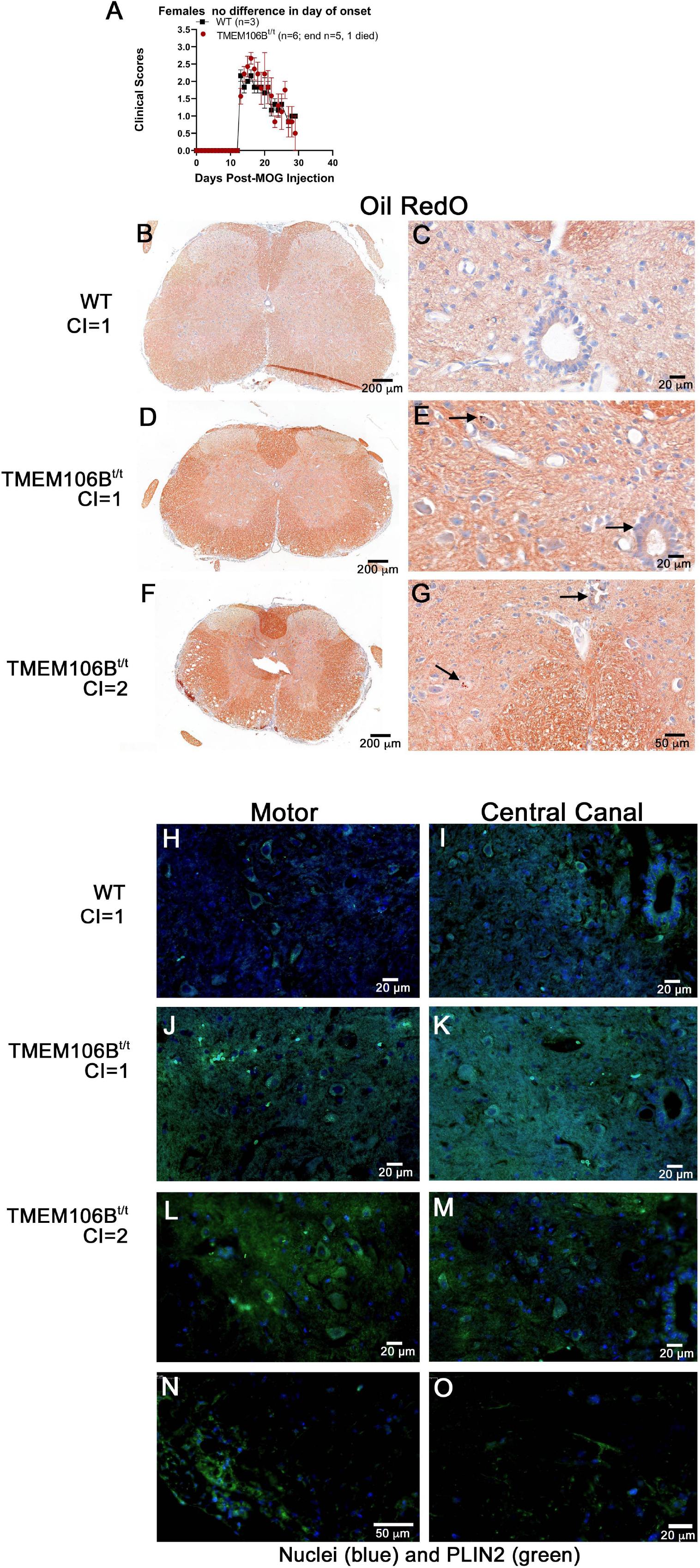
TMEM106B^t/t^ mice have significant OilRedO+ deposition in the corpus callosum at 2- and 3weeks post-cuprizone withdrawal. **(A,B)** illustrates the lack of OilRedO+ staining in the corpus callosum of control, naïve WT and TMEM106B^t/t^ mice at 5.5 months of age. (**C-J**) 10-week old mice were given a 5-week cuprizone diet, followed by regular chow for 2-weeks (5+2; **C,D,G,H**) and 3-weeks (5+3; **E,F,I,J**) prior to sacrifice. **(G-J)** A myelin stain (Brain Stain Imaging kit B34650; Molecular Probes) shows no significant change in remyelination at 2- weeks (**G,H**) and 3-weeks (**I,J**) post-cuprizone withdrawal. (**K**) Mann-Whitney U test determined no difference in Iba1 (4.5, >0.9999), SMI32 (4.5, p=>0.9999), APC/CC1 (0.5000, p=0.2000) and CAII (2, p=0.4000) immunopositive cells/mm^2^ at 2-weeks post-cuprizone withdrawal. Mann-Whitney U test shows no difference in SMI32 (**L**), APC/CC1 and CAII immunopositive cells/mm^2^ (**M**) or myelin (**N**) mean fluorescent intensity (4, p=0.6286) at 3-weeks post-cuprizone withdrawal. WT (n=4), TMEM106B^t/t^ (n=3).

## Discussion

We have identified the enrichment of TMEM106B in β-mercaptoethanol/sarkosyl-insoluble pellets from white matter plaques isolated from individuals with RRMS by mass spectrometry, and observe increased TMEM106B immunoreactive puncta in human white matter from individuals with RRMS relative to Alzheimer’s disease and non-neurologic controls. *In vitro* and *in vivo* studies show that TMEM106B protein levels are known to affect lysosomal function.^12,22^ TMEM106B is observed in dendrites and axons, particularly in the axonal initial segment where it is suggested to mediate axonal sorting in facial nerve and sorting of lysosomes in axons, where vacuolization was detected predominantly in motor neurons of TMEM106B knockout mice.^4,9^ We observed axons with TMEM106B staining in the cortex of individuals with RRMS, Alzheimer’s disease and non-neurological controls, its normal localization. In contrast, low TMEM106B levels result in clustering of small lysosomes near the nucleus; and in neurons, altered retrograde lysosomal mobility and shortened dendritic branching were reported.^10^ ‘High levels of TMEM106B result in larger and fewer lysosomes that fail to produce normal intra-lysosomal acidity, resulting in lysosomal stress.^22^ The presence of TMEM106B in sarcosyl-insoluble pellets from the plaques of individuals with MS suggests that sequestration of TMEME106B within insoluble pellets may result in lysosomal dysfunction/stress perhaps with delayed and inefficient clearance of debris within the plaque. OilRedO+ neutral lipid droplets stain myelin debris within and adjacent to MS plaques,^23–26^ with deposition reported within macrophages/microglia, and not within astrocytes.^25,26^ We observed OilRedO+ deposits in frozen sections of MS plaques from the individuals in which we performed nanoLC-ms/ms, although we did not observe TMEM106B+/OilRedO+ cells within all frozen MS brain sections examined (data not shown).

Post-mortem material is static, and does not allow for the type of analysis mouse models permit. The TMEM106B^t/t^ mouse allowed us to examine the impact of impaired TMEM106B in two models of nervous system injury, MOG-induced EAE and cuprizone-induced demyelination and remyelination. While our prediction was that the TMEM106^t/t^ mice would have higher clinical scores than WT mice during acute and chronic EAE, we observed the same clinical course in both groups. However, during chronic EAE we observed increased OilRedO+/PLIN2+ lipid deposits in TMEM106B^t/t^ spinal cords suggesting that there is lipid droplet accumulation and likely a failure to efficiently clear debris following injury, likely contributing to a delay in recovery. The mere association of PLIN2 with lipid droplets suggests lipophagic activity. In addition, we observed more axonal damage, reflected in an increase in SMI32+ axonal swellings and less intact myelin, both by a myelin stain and MBP immunostaining during chronic EAE. Our TMEM106B^t/t^ mice are hypomorphic and express low levels of TMEM106B, and relatively normal levels of myelin proteins during development^11^. A TMEM106B CRISPR/Cas knockout mice have reduced myelination and less PLP at the membrane during oligodendrocyte development.^13^

Toxic fatty acids can accumulate within neurons due to a failure of lipid particles to transfer the fatty acids to astrocytes where they are detoxified.^27^ The ability to detect OilRedO+/PLIN2+ lipid droplets within the cell bodies of TMEM106B^t/t^ mice suggests that lipid droplet accumulation contributes to neuronal damage and loss during EAE. PLIN2 is a lipid droplet resident protein associated with the metabolism of intracellular lipid droplets, and PLIN2 becomes more stable when associated with lipid droplets.^28^ PLIN2 and PLIN3 expression are ubiquitous, with PLIN1 mainly in adipose tissue.^29^ Indeed, we determined that there was a correlative increase in OilRedO+ and PLIN2+ lipid droplets in TMEM106B^t/t^ spinal cords during chronic EAE. Future studies will evaluate changes in lipid profiles in spinal cords of WT and TMEM106B^t/t^ mice during chronic EAE by lipidomics.

TMEM106B is a type II transmembrane glycoprotein that localizes to lysosomes.^6,30,31^ With limited data on TMEM106B’s normal function *in vivo,* this study and several other groups have begun to further explore the function of TMEM106B in the CNS using TMEM106B^-/-^ mice.^3,11,13,14,22,32^ Beneficial effects of reduced TMEM106B levels have been reported, especially in frontotemporal dementia-related phenotypes in progranulin-deficient mice where the loss of TMEM106B ameliorated abnormal lysosomal phenotypes.^16^ However, TMEM106B has opposing effects in different mouse models of lysosomal diseases when Gaucher disease and disease and neuronal ceroid lipofuscinosis were examined. In cultured primary hippocampal neurons, knockdown of TMEM106B affects the transport of lysosomes in dendrites, ultimately leading to reduced dendritic branching. ^9,10^ The knockdown of TMEM106B in HeLa cells resulted in the redistribution of lysosomes from the cell periphery to the perinuclear region^10^ and a reduced number of lysosomes per cell.^9^ In facial motor neurons, proximal axonal swelling with enlarged LAMP1+ vacuoles and an accumulation of autophagolysosomes revealed that TMEM106B functions in axonal transport of LAMP1+ organelles and axonal sorting at the initial segment.^4^

Our TMEM106B^t/t^ mouse model retains low levels of TMEM106B protein by western blot analysis.^9,11^ This reduced protein level is insufficient to maintain homeostasis within the CNS and efficiently clear lipid-laden debris in challenged mice. Of the CRISPR/Cas mouse models that generated total knockout of TMEM106B^5,9,16,32^, two independent groups showed loss of TMEM106B led to myelination deficits.^3,12,13^ While our cuprizone studies with TMEM106B^t/t^ mice were in progress, CRISPR/Cas TMEM106B knockout mice were generated showing changes in myelin following cuprizone-induced demyelination/remyelination.^13^ Similar to our study, they found no significant difference in the number of oligodendrocytes within the corpus callosum during recovery, or a significant change in myelin. They did not examine or report persistent myelin debris with increased OilRedO+ deposition during 2- and 3-weeks recovery from cuprizone-induce demyelination. Combined studies suggests that during an initial recovery from cuprizone-induced toxicity, oligodendrocytes have the capacity to remyelinate even in the presence of myelin debris. Whether there is decreased ability to remyelinate following multiple episodes of demyelination/remyelination similar to what is observed in MS needs to be tested. Future studies will determine whether prolonged cuprizone treatment (12-weeks), or a second 5-week cuprizone treatment course after the initial treatment and recovery results in deficits in remyelination in TMEM106B^t/t^ treated relative to control mice. In addition, the cuprizone model in combination with EAE can be performed.^33^ Our combined data demonstrate that in mice, reduction in TMEM106B in the spinal cord during EAE has a more profound effect than in the corpus callosum, where remyelination can occur in the presence of OilRedO+ deposition. The presence of OilRedO+ neutral lipids within the CNS suggest that TMEM106B has a functional role in the clearance of lipids by a specialized form of autophagy, known as lipophagy. Lipophagy functions to regulate intracellular lipid stores, cellular levels of free lipids such as fatty acids and energy homeostasis.^34^ A recently published manuscript detailed structural similarities between human TMEM106B and two yeast proteins Vac7 and Tag1, predicting that all three proteins have LEA-2 domains and are lipid transfer proteins.^35^ The C-terminus of TMEM106B and LEA-2 are highly homologous. Vac7, a regulator of PI(3,5)P2 stress responses and production, has one LEA-2 domain. Tag1, which signals to terminate autophagy, has three LEA-2 domains, and TMEM106B has one LEA-2 domain. The author speculates that these proteins may sense a signal derived from autophagic material build-up, possibly lipids and communicate to regulate autophagy. In conclusion, TMEM106B has the potential to regulate lysosomal function, and when TMEM106B levels are reduced there is retention of OilRedO+/PLIN2+ lipids following CNS injury that contributes to CNS impairment. Lipid droplet accumulation was observed in TMEM106B^t/t^ mice under conditions where WT and TMEM106B^t/t^ mice had identical clinical scores during recovery and chronic EAE, and at the same times during recovery from cuprizone-induced demyelination. The notion that TMEM106B may sense a signal derived from lipids and communicate signals to regulate autophagy is intriguing and warrants further examination.

Selective autophagy is important for sequestering protein cargo into autophagosomes and targeting autophagy receptors to LC3 (microtubule-associated proteins 1A/1B light chain 3B) for delivery to the lysosome. Several proteins containing LC3 interacting region (LIR)-motifs have been identified,^36^ and we examined the proteins in our RRMS insoluble pellet and identified several unique LIR-peptides within the aggregate including perhaps TMEM106B. TMEM106B contains a possible LC3 LIR motif [Y/F/W]X1-X2[L/I/V] at amino acid 18-21, in a natural disordered region EDAYDGVTSE (PONDR.org, Figure 8). The **LIR** is often flanked by diverse sequences containing Ser, Thr and/or the negatively charged residues Glu (E) and/or Asp (D). This region is seen in proteins that function in macroautophagy and clearance of debris.^36^ Although an *in vitro* study determined that mutation of the tyrosine within the consensus LIR motif still allowed TMEM106B to localize to the lysosome.^37^ Several proteins have been suggested to bind within TMEM106B’s disordered regions^38^ and future studies will test whether the alteration in the motifs limit efficient clearance of the insoluble proteins and lipids. Alterations in TMEM106B function have been linked to hypomyelinating leukodystrophy,^39^ Pick’s disease, frontotemporal dementia with TDP-43 inclusions^40^; age-related TDP-43 encephalopathy (LATE), Parkinson’s disease, and defects in dendritic trafficking of lysosomes^10,41–43^ and axons.^4^ The presence of TMEM106B within β-mercaptoethanol/sarkosyl-insoluble pellets following a high-speed ultracentrifugation suggests that a significant proportion of TMEM106B in MS plaques is unavailable to traffic to the lysosome to carry out its function in clearance. Dysfunction of TMEM106B is known to impair formation, transport and function of lysosomes^44^ that can have profound effects on recovery from demyelinating lesions in RRMS.

**Figure 8:**
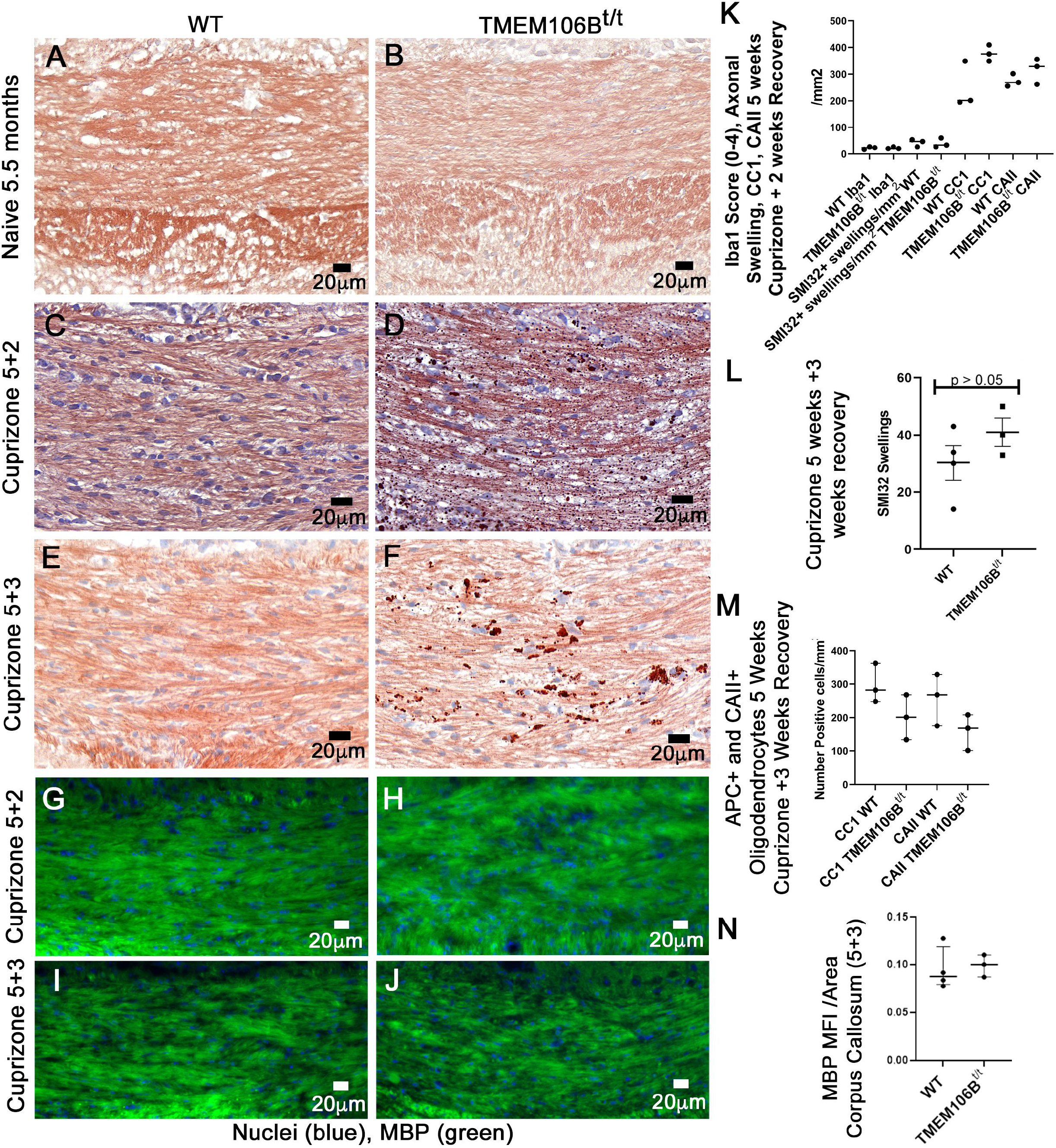
Regions of disorder and a possible LIR motif [W/F/Y]-X1-X2-[L/I/V] for potential lysosomal localization within TMEM106B protein (pndr.com).

## Acknowledgements

We acknowledge the following sources of human tissue used in this study: The Rocky Mountain MS Center at the University of Colorado, and its grant support from the National Multiple Sclerosis Society (NMSS # SI-2011-37165); the UCLA Brain Bank and its NMSSS grant support G-1510-06784; and Dr. Steven Chin, Director of Neuropathology, Dr. Kathleen Whitney, Dr. Anita Arackal, and Dr. Ukuemi Edema, Department of Pathology, Montefiore Medical Center, and The Montefiore Bank Repository for help identifying and sectioning additional control brain specimens. We are grateful to Ms. Candace Bretsch, RMMSC Tissue Bank Coordinator, who was exceptionallly helpful in providing information for the redacted samples used in the study. We thank Dr. Cristina C. Clement, Weill Cornell Medical Center for Immunology, Radiation Oncology, for providing the protocol for the isolation of the insoluble proteins, and advice on mouse aggregate studies prior to initiation of the human studies. We thank Dr. Rajat Singh for discussions on LIR-domains, and Mr. Eddie Nieves for helpful discussions on mass spectrometry and proteomics. We thank Dr. Yongtai Lo for advice on statistical analysis of the human samples following mass spectrometry, and Ms. Andrea Briceno for scanning stained slides. We thank Susmita Kaushik for providing PLIN2 antibody. We thank Dr. Jeffrey Bonanno for helping with pondrfit analysis of TMEM106B. We thank Mr. Emilio Merheb for reading and commenting on the manuscript.

## Abbreviations

APC: adenomatous polyposis coli.
CAII: carbonic anhydrase II.
CI: clinical index.
EAE: experimental autoimmune encephalomyelitis.
GFAP: glial fibrillary acid protein.
Iba1: Ionized calcium binding adaptor molecule 1.
PLIN2: perlipin2.
RRMS: relapsing remitting multiple sclerosis.
SPMS: secondary progressive multiple sclerosis.
MOG: myelin-oligodendrocyte glycoprotein.
MBP: myelin basic protein.
MS: multiple sclerosis.
TMEM106B: transmembrane protein106B.
WT: wildtype.

---

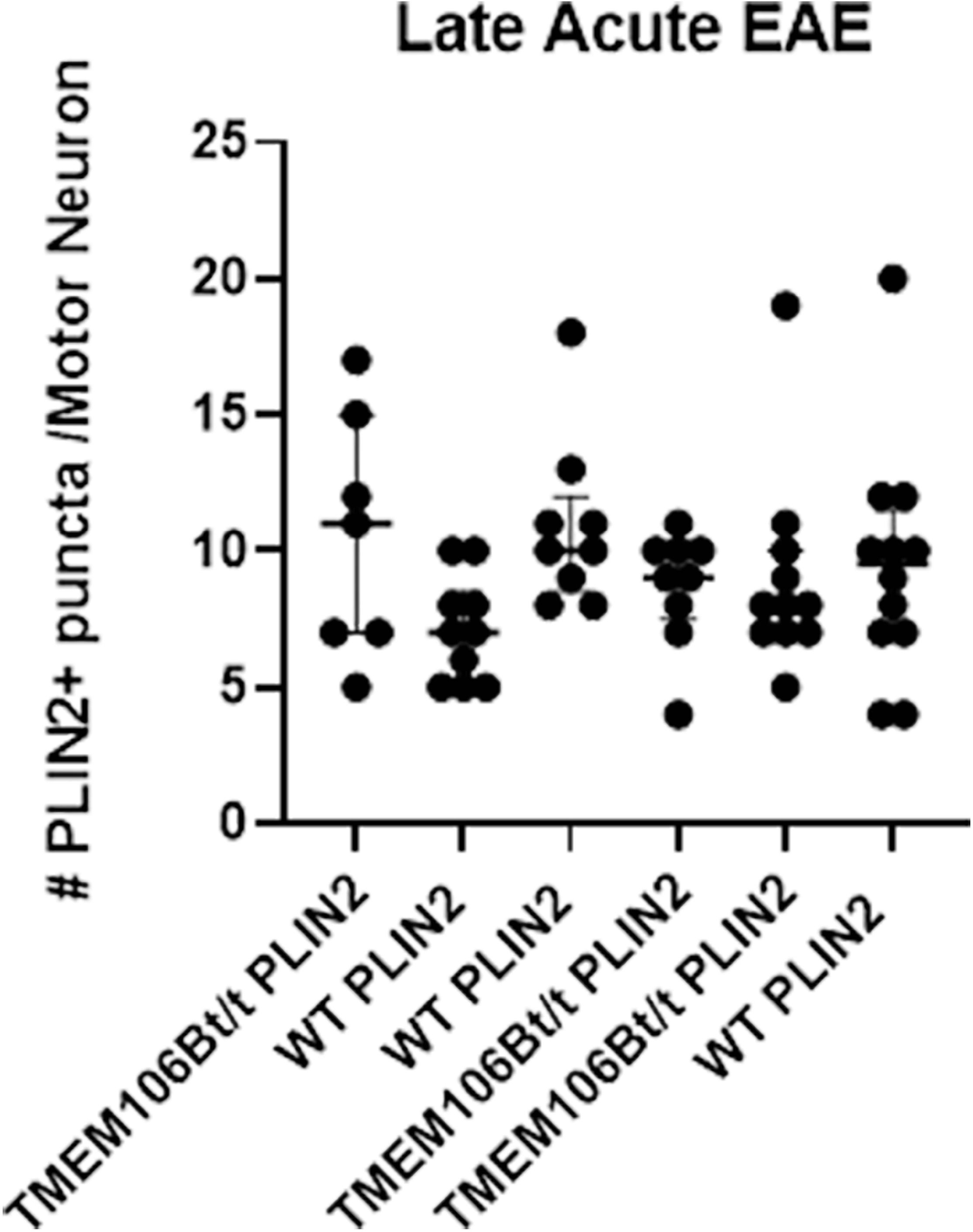

**Supplemental Figure 1:**
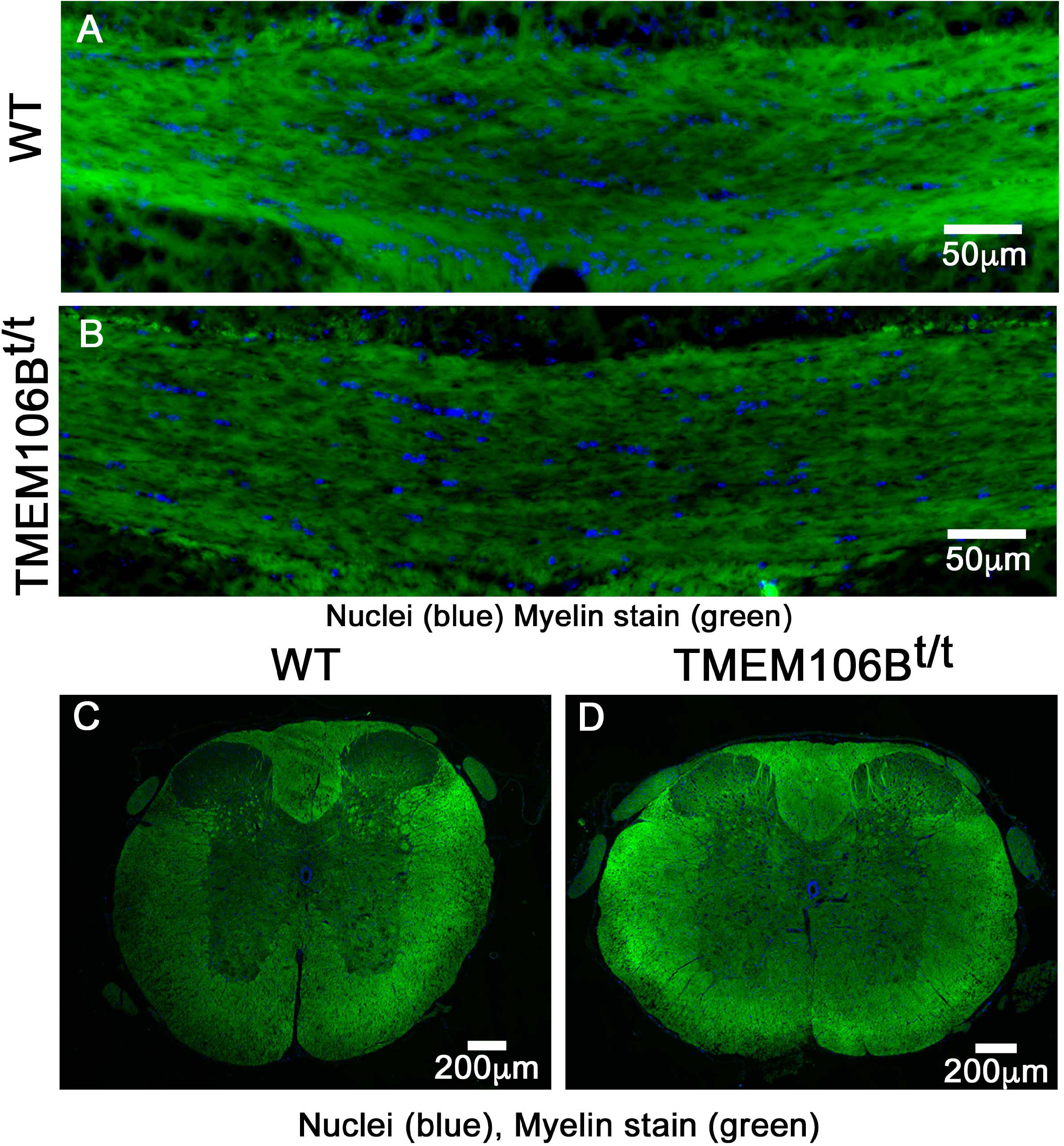
Naïve WT and TMEM106B^t/t^ mice have similar myelin deposition in brain (A,B) and spinal cord (C,D). Naïve mice at 10 weeks of age were stained with the fluoromyelin and DAPI components of the BrainStain kit (Invitrogen).

**Supplemental TABLE 1:** Available samples for the ongoing study.

**RRMS**
| Case # | PMI* | Age | Gender | Diagnosis |
| --- | --- | --- | --- | --- |
| 1 | 5h | 65 | Female | RRMS |
| 2 | 2h | 69 | Female | RRMS |
| 3 | 7h | 64 | Female | RRMS |
| 4 | 5h | 65 | Female | RRMS |
| 5 | 24h | 61 | Male | RRMS |

\*Post-mortem interval

**Non-neurological White Matter Controls**
| Case # | PMI | Age | Gender | Diagnosis |
| --- | --- | --- | --- | --- |
| 6 | 5h | 57 | Female | Breast Cancer |
| 7 | 5h | 74 | Male | Cardiac |
| 8 | 9h | 63 | Male | Infarct/Cardiac |
| 9 | 19h | 59 | Male | Cardiac |
| 10 | 6.5h | 65 | Male | Cardiac |

**For IF and IHC Staining:** **MS Cases Paraffin-embedded**
| Case | PMI | Age | Gender | Diagnosis |
| --- | --- | --- | --- | --- |
| 11 | 15h | 49 | Female | Secondary Progressive, Plaque and NAWM |
| 12 | <10h | 61 | Female | RRMS, Plaque and NAWM |
| 13 | 10h | 61 | Male | RRMS, Plaque and NAWM |
| 14 | UNK | 68 | Male | MS type unknown, Plaque and NAWM |
| 15 | 14h | 73 | Male | RRMS, Plaque and NAWM |
| 16 | 21h | 65 | Female | RRMS, Plaque and NAWM |
| 17 | 14h | 64 | Male | RRMS, Plaque and NAWM |
| 18 | 12h | 69 | Male | RRMS, Plaque and NAWM |
| 19 | 12h | 57 | Male | RRMS, Plaque and NAWM |
| 20 | 18h | 68 | Male | RRMS, Plaque and NAWM |

**Other Neurologic Controls and Non-Neurologic Controls** **Alzheimer's Disease Paraffin-embedded**
| Case # | PMI | Age | Gender | Diagnosis |
| --- | --- | --- | --- | --- |
| 21 | UNK* | 86 | Female | AD |
| 22 | UNK | 66 | Female | AD |
| 23 | UNK | 62 | Male | AD |
| 24 | 12 h | 70 | Male | AD |
| 25 | UNK | 63 | Male | AD |
| 26 | UNK | 80 | Female | AD |
| 27 | UNK | 61 | Male | AD |

\*Unknown

**Non-Neurological Controls (NNCs) Paraffin-embedded**
| Case # | PMI | Age | Gender | Diagnosis |
| --- | --- | --- | --- | --- |
| 28 | 2 d | 72 | Male | Pneumonia |
| 29 | 22h | 65 | Female | Septic shock |
| 30 | UNK | 78 | Male | No diagnostic abnormality recognized |
| 31 | UNK | 67 | Male | No diagnostic abnormality recognized |
| 32 | UNK | 62 | Male | No diagnostic abnormality recognized |
| 33 | UNK | 81 | Male | Primary Age-related tauopathy |
| 34 | UNK | 26 | Male | Severe and chronic lung injury, respiratory distress syndrome, liver cirrhosis |
| 35 | UNK | 59 | Male | Advanced K-ras-mutated non-small cell adenocarcinoma stage IV |
| 36 | UNK | 48 | Female | Metastatic carcinoma probable urachal origin and hypertension |
| 37 | UNK | 68 | Female | Toxic epidermal necrolysis 60% body |

